## Supplementary Method for "Planar cell polarity coordination in a cnidarian embryo provides clues to animal body axis evolution"

### List of solution and concentrations

#### Morpholino oligos

| Name | target gene | Genbank | marimba acc. | Morpholino seqeunce | Blocking action | Concentration |
|---|---|---|---|---|---|---|
| Wnt3MO | CheWnt3 | EU374721.1 | XLOC_001931 | CCAAAACACACCAGTGTCGAGCCAT | translation | 0.1 mM |
| Fmi-MO | CheFmi | JQ439002.1 | XLOC_001639 | CCTCAAGCCATCTGAGCTTCATTTT | translation | 2 mM |
| Stbm-MO | CheStbm | JQ439008.1 | XLOC_013328 | TCACTCCATCATCAAAATCATCCAT | translation | 0.32 mM |
| Fat-L-MO | CheFat1 | JQ439001.1 | XLOC_033731 | CATCAAAATGTGAGACTTACCAGCC | splicing | 0.5 mM |
| Ds-MO | CheDs | JQ438999.1 | XLOC_037142 | TACGGTGATGGATAGTTCATCTTTC | translation | 4 mM |

#### Dye

| Name | Source | Catalog number | Cocnentration |
|---|---|---|---|
| Phalloidin Alexa Fluor™ 488 | Invitrogen | A12379 | 4 unit/ml 0.13 μM in PBS |
| Phalloidin Alexa Fluor™ 647 | Invitrogen | A22287 | 4 unit/ml 0.13 μM in PBS |

#### Antibodies

| Name | Source | Catalog number | Host |
|---|---|---|---|
| anti-Par3 | Merk Millipore | 07-330 | rabbit polyclonal |
| anti-γ Tubulin GTU-88 | SIGMA | T5326 | mouse monoclonal |
| anti-centrin 20H5 | Merck Millipore | 04-1624 | mouse monoclonal |
| Rhodamine Red-AffiniPure Anti-Mouse IgG (H+L) | Jackson immunoResearch | 115-295-003 | Goat |
| Alexa Fluor 555 AffiniPure Goat Anti-Rabbit IgG (H+L) | Jackson immunoResearch | 111-565-144 | Goat |

#### Synthetic mRNA

| Name | Gene | species | Genbank | Marimba acc. | Role | Modification | conc ng/μl |
|---|---|---|---|---|---|---|---|
| Wnt3 | CheWnt3 | Clytia | EU374721.1 | XLOC_001931 | Wildtype | 3 bp mismatch in MO target | 100 |
| dnTCF | CheTCF | Clytia | JQ806376.1 | XLOC_007658 | Dominant ngegative | N-terminal 27 a.a. deletion | 400 |
| CA-β-cat | Che-β-catenin | Clytia | JQ438997.1 | XLOC_003560 | Constitutively active | S97A, T101A, S105A | 20 |
| RhoABC-DN | CheRhoABC | Clytia | | XLOC_044216 | Dominant ngegative | T19N | 10 |
| RhoABC-CA1 | CheRhoABC | Clytia | | XLOC_044216 | Constitutively active | G14V | 10 |
| RhoABC-CA2 | CheRhoABC | Clytia | | XLOC_044216 | Constitutively active | Q63L | 10 |
| PH-mCherry | Phosphoinositide phospholipase C-delta-1 | Human | U09117.1 | | Label membrane | Codon-optimised PH-domain (N-terminal 174 aa) with mCherry at C-terminal | 100 |
| Poc1-Venus | ChePoc1 | Clytia | HM010924.1 | XLOC_001641 | Label centriole | full length ChePoc1 + Venus codon optimised | 2 |

http://marimba.obs-vlfr.fr

#### PCR primers for mRNA template

| Name | Sequence (5'-3') | binds |
|---|---|---|
| T3-PH-F | GCAATTAACCCTCACTAAAGGGTAAGAAAACAAAGTAAACAAAC ATGGATAGCGGGCGAGATTTTTTAAC | PH-domain 5' |
| T3-Poc1-F | GCAATTAACCCTCACTAAAGGGTAAGAAAACAAAGTAAACAAAC ATGACTTCTGCTTTGGAGGATCC | Poc1 5' |
| CC-UTR-R | TTTATTAAGCCTAAAATAAATTACTTACTTA TTATTTATAAAGCTCATCCATACCACCCG | mCherry (codon optimized) 3' |
| Venus-UTR-R | TTTATTAAGCCTAAAATAAATTACTTACTTA TTACTTGTACAGCTCGTCCATGCC | Venus 3' |
| CC-UTR-R | TTTATTAAGCCTAAAATAAATTACTTACTTA TTATTTGTAAAGTTCGTCCATACCAAGAGTT | Venus (codon optimized) 3' |

### Image segmentation

1) Open confocal scan image with ImageJ (Fiji).
2) Determine the Z-axis range and  Z-project the images.
   - Image/Stacks/Z project… (Max Intensity)

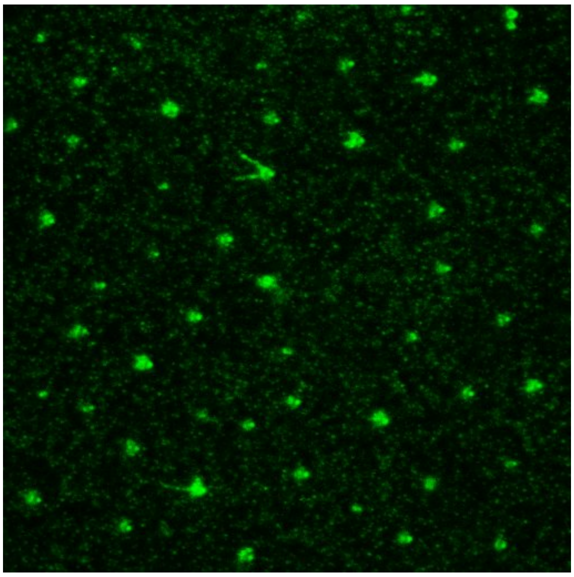

γ-tubulin

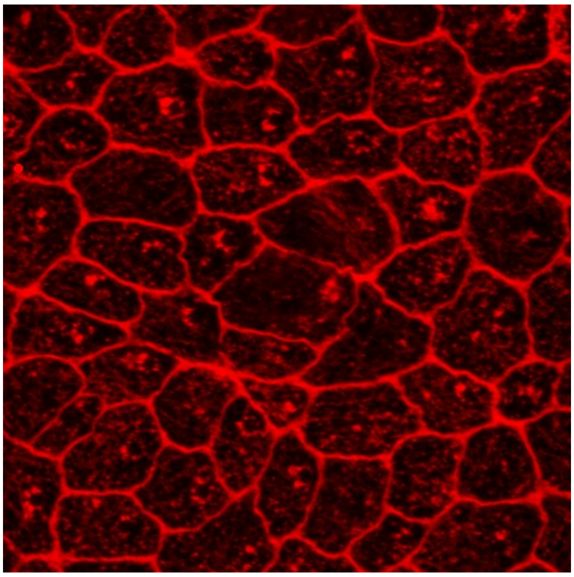

actin

3) Split channels.
4) Manually track the basal body and cell contours by Pencil Tool (with a width 3~4-pixel for actin (0.2~0.3 µm) and up to 10-pixel size for the basal body (~0.8 µm diameter)
   - For the basal body, pick between two dots, most likely corresponding to two centrioles.

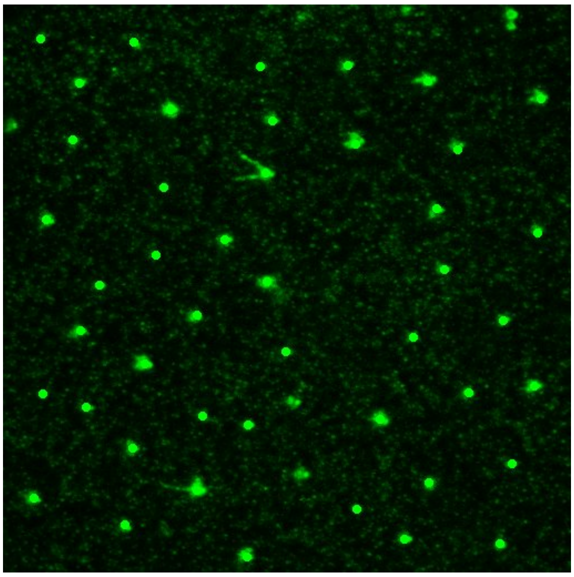

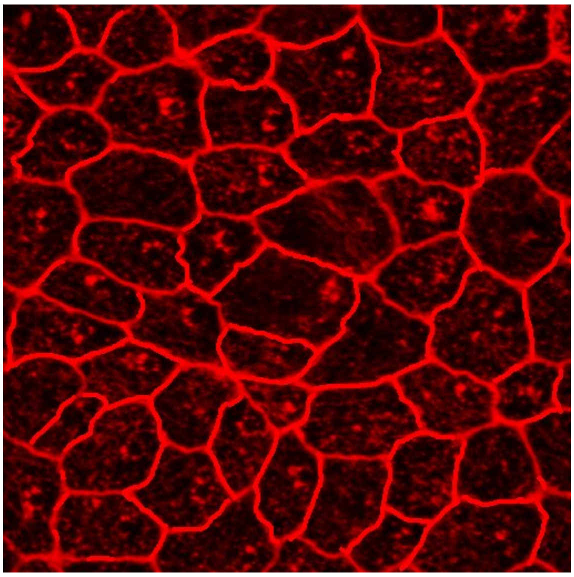

5) Define the threshold for binary image conversion. (Image/Adjust/Threshold) then convert to binary images (Process/Binary/Make Binary). If necessary, invert the image so that the background becomes black.

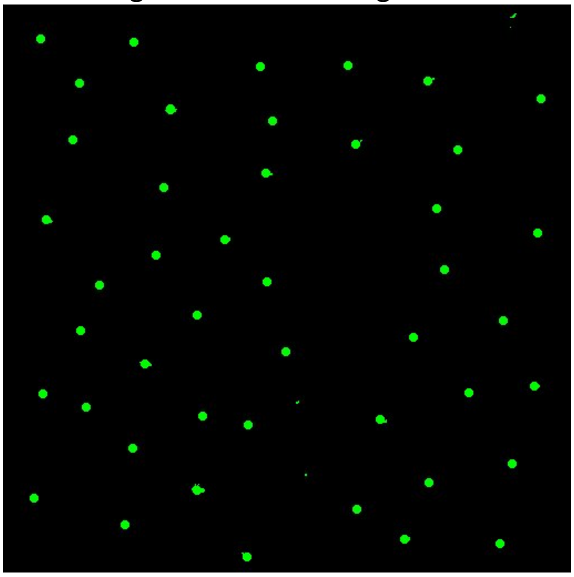

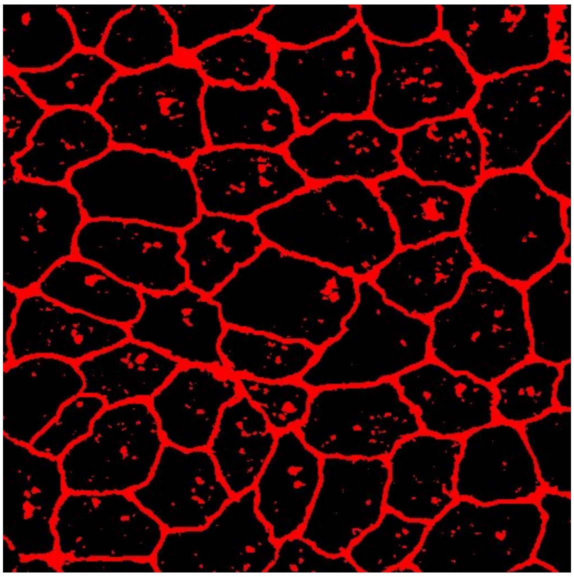

6) For the actin channel, apply binary segmentation from Process/Binary menu
   a. Close –
   b. Erode
   c. Dilate
   d. Repeat b and c

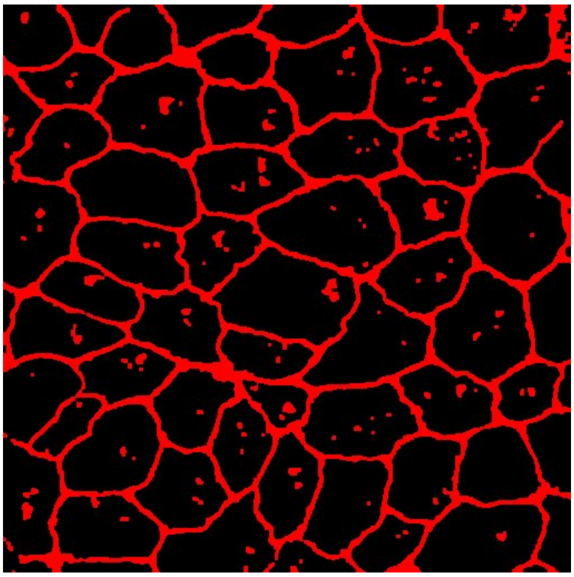

7) Set measurement parameters (Analyze/Set Measurements…)
   a. Select Area, Centroid (it will be the geographical center of cell) Center of Mass (it will be basal body position)
   b. Select the image of γ-tubulin channel for "Redirect to" option
8) Measure (Analyze/Analyze Particles)
- Size (specify 2-Infinity, micron^2)
- Show: Outlines
- Check "Display results", "Exclude on edges" and "Clear results".

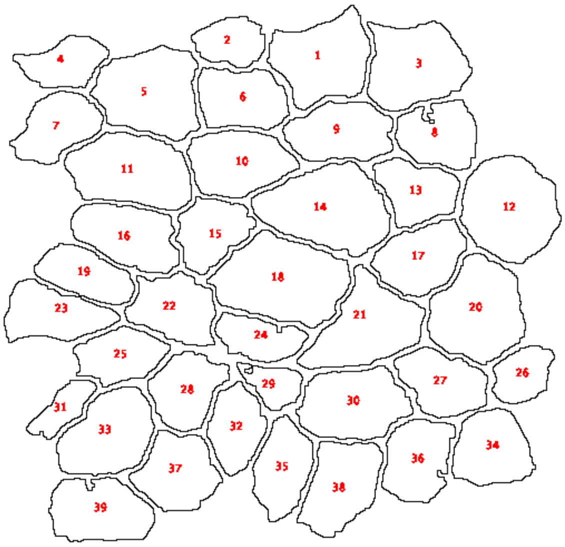


| | Area | X | Y | XM | YM |
|---|---|---|---|---|---|
| 1 | 34.130 | 23.376 | 5.176 | 23.737 | 4.095 |
| 2 | 14.047 | 16.793 | 3.913 | 18.500 | 4.405 |
| 3 | 34.605 | 30.000 | 5.622 | 30.640 | 5.432 |
| 4 | 16.023 | 4.830 | 5.431 | 5.446 | 5.606 |
| 5 | 37.382 | 10.987 | 7.566 | 12.126 | 7.495 |
| 6 | 23.341 | 17.513 | 8.008 | 19.381 | 8.329 |
| 7 | 21.577 | 4.293 | 10.037 | 4.965 | 9.691 |
| 8 | 21.218 | 31.743 | 10.662 | 32.756 | 10.412 |
| 9 | 26.099 | 24.664 | 10.450 | 25.409 | 9.997 |
| 10 | 27.138 | 17.259 | 12.612 | 18.965 | 12.133 |
| 11 | 37.927 | 9.623 | 13.237 | 11.533 | 13.135 |
| 12 | 41.064 | 36.839 | 15.918 | 38.523 | 16.418 |
| 13 | 18.883 | 30.086 | 14.631 | 31.355 | 14.616 |
| 14 | 43.770 | 23.750 | 15.923 | 23.750 | 15.923 |
| 15 | 19.531 | 15.409 | 17.616 | 15.986 | 16.900 |
| 16 | 27.382 | 9.371 | 17.870 | 10.972 | 18.020 |
| 17 | 24.528 | 30.426 | 19.485 | 31.795 | 19.061 |
| 18 | 45.605 | 20.404 | 21.116 | 18.981 | 19.942 |
| 19 | 18.736 | 6.321 | 20.706 | 6.888 | 20.182 |
| 20 | 38.395 | 34.551 | 23.340 | 36.040 | 22.745 |
| 21 | 35.470 | 26.518 | 24.461 | 29.553 | 23.946 |

9) Check separation of individual cells. If necessary repeat 3)~7)
   (X, Y) is the coordinate of the centroid of the apical surface (**$x_c$, $y_c$**), and (XM, YM) is the coordinate of the basal body (**$x_b$, $y_b$**).
   "Area" column measures the apical surface area **S** in μm$^2$
10) Save the result (csv format).

### PCP statistics

In MATLAB, import the file and analyse it with `plotpcp.m` script (below.)
Example

```
pcpplot(xc, yc, xb, yb, 246, 'wnt3rescue')
```

where 4$^{th}$ argument is the width/height of the field of view (246 μm), 5$^{th}$ argument is prefix for output files.

### Polarisation index

The polarization index is defined as the distance between the centroid of the apical surface and the basal body relative to the circle's radius with the same surface.

$$\frac{\sqrt{(x_c - x_b)^2 + (y_c - y_b)^2}}{\sqrt{\frac{S}{\pi}}}$$

```
function plotpcp(x1, y1, x2, y2, varargin)
%% common data treatment

% (x1, y1): basal body
% (x2, y2): cell cenroid
% VARARGIN:
%       'label', label for outpout
%       'fov', length of the longer edge of the field of view rectangle
%       in µm, default 246 µm
%       'revy', 1 or 0, reverse orientation y axis if 1
%       'color', color map default hsv
%       'factor', conversion factor (if input unit is not µm)
%       'separation', minimum seperaton of cell centroid and basal body
%       position to consider the cell is polarized, default 0.5 µm

% important note: this script assumes the origin of x-y coordinate is top
% left, adapting to the ImageJ "measurement" fonction.

if nargin < 4
  error('x1, y1, x2 and y2 are needed (x1,y1):basal body (x2, y2):centroid');
  return
end

fov=246.0;
label = 'result';
revy = 1;
cmap = 'hsv';
factor = 1.0 ; % scaling factor
limit = 0.5 * 2; %minimum polarization (distance between bb and centroid x 2)



for i=1:length(varargin)
  vin = varargin{i};
  if isequal(vin,'label')
    label = varargin{i+1};
  end
  if isequal(vin,'fov')
    fov = varargin{i+1};
  end
```

```
  if isequal(vin,'revy')
    revy = varargin{i+1};
  end
  if isequal(vin,'color')
    cmap = varargin{i+1};
  end
  if isequal(vin,'factor')
    factor = varargin{i+1};
  end
  if isequal(vin,'separation')
    limit = varargin{i+1} * 2;
  end

end

x1 = x1 * factor;
y1 = y1 * factor;
x2 = x2 * factor;
y2 = y2 * factor;
u = (x2-x1)*2; % vector
v = (y2-y1)*2; % vector

plotboundary = ((-12:12) + 0.5 )* pi /12;% 24 categories
tickboundary = ((0:11) + 0.5 )* 30;
ticklabel = [ " ", "Wnt3(+) side" " " " " "90" " " " " "180" " " " " "-90"  ];


%% orientation and length

orientation = atan2(v,0-u);  % AAN2 (Y-vector, X-vector)
% given top left corner as the origin
% south (0,1) -> 0
% north (0,-1) -> 3.14
% east (1, 0) -> 3.14/2
% west (-1, 0)  -> -3.14/2

vec_len = vecnorm([u, v],2,2); % vnorm is length of the vector

% remove non-polarized cells and register
orientation(vec_len <= limit) = NaN;


% non-polarized subset
unpolx = x2(isnan(orientation));
unpoly = y2(isnan(orientation));

% polarized subset
```

```
x1pol = x1(~isnan(orientation)); % x1pol is x1 for only polarized cells
x2pol = x2(~isnan(orientation));
y1pol = y1(~isnan(orientation));
y2pol = y2(~isnan(orientation));
upol = (x2pol-x1pol)*2; % vector x
vpol = (y2pol-y1pol)*2; % vector y

orientationpol = rmmissing(orientation);  % orientationpol is only polarized cells

colorvalue = mod(- orientationpol/(2*pi), 1) ; % adjusting color range from 0 to 1



%% histogram of PCP orienttion

% rador plot

f_rador = figure;

polarhistogram(orientationpol, plotboundary ,'Normalization', 'probability');

ax = gca;

set(ax,'ThetaTick', tickboundary, 'ThetaTickLabel', ticklabel);
set(ax,'FontSize',8);
set(ax, 'RAxisLocation', 120);

outerpos = ax.OuterPosition;
ti = ax.TightInset;

left = outerpos(1) + ti(1);
bottom = outerpos(2) + ti(2);
ax_width = outerpos(3) - ti(1) - ti(3);
ax_height = outerpos(4) - ti(2) - ti(4);
ax.Position = [left bottom ax_width ax_height];

set(f_rador, 'units','centimeters','position',[35, 40, 4 , 4])

savefig(f_rador, strcat(label, '_h-ori', '.fig'));


%% histgram of basal body - centroid distance
f_len = figure;
histogram(vec_len, 40);
savefig(f_len, strcat(label, '_h-len', '.fig'));
```

```
%% PCP vector plot
f_plot = figure; % make a new figure

% polarized cells (bar)
handle = vfield(x1pol, y1pol, upol, vpol, colorvalue,'tr', 0);% vector plot the color is specified
by orientation3
xticks([]);
yticks([]);
set(handle, 'LineWidth', 2.75);
pbaspect([1 1 1]);
box on;

colormap(cmap); % This should be changed to your favorite color
caxis([0 1]) % caxis fix minimum and maxium value for the left and right-end of the color bar

% non-polarized cells (dot)
hold on; % hold on to "overlay" 2nd plot on the existing plot
scatter(unpolx, unpoly, 14, 'black', 'filled'); % scatter plot of the non-polarized cells

ax = gca;
if revy==1
    set(ax ,'YDir','reverse'); % reverse Y-axis so that top left will be the origin
end
xlim([0 fov]); % fix the range of the plot
ylim([0 fov]);

pbaspect([1 1 1]); % fix aspect ratio
hold off;

savefig(f_plot, strcat(label, '_plot', '.fig'));


%% some statistics
num_of_total_cells = numel(x1);
num_of_non_polarized_cells = numel(unpolx);
num_of_polarized_cells = numel(x1pol);
non_polarized = num_of_non_polarized_cells / num_of_total_cells * 100;

mean_orientation= circ_mean(orientationpol) *180/pi; % in degree
[ang_dev, cir_std ]= circ_std(orientationpol); % *2 *180 / pi % 0.95 confidence interbval

ang_dev = ang_dev *180 / pi;
cir_std = cir_std *180 / pi;

fprintf("%d cells polarized, %d non polarized out of %d cells. (%.1f parcent non
polarized)\n", num_of_polarized_cells, num_of_non_polarized_cells, num_of_total_cells,
non_polarized);
```

```
fprintf("mean orientation %.1f° circular standard deviation %.1f°\n", mean_orientation,
cir_std);
```
