## Supplementary Figures 1-6 for "Planar cell polarity coordination in a cnidarian embryo provides clues to animal body axis evolution"

**Figure S1**

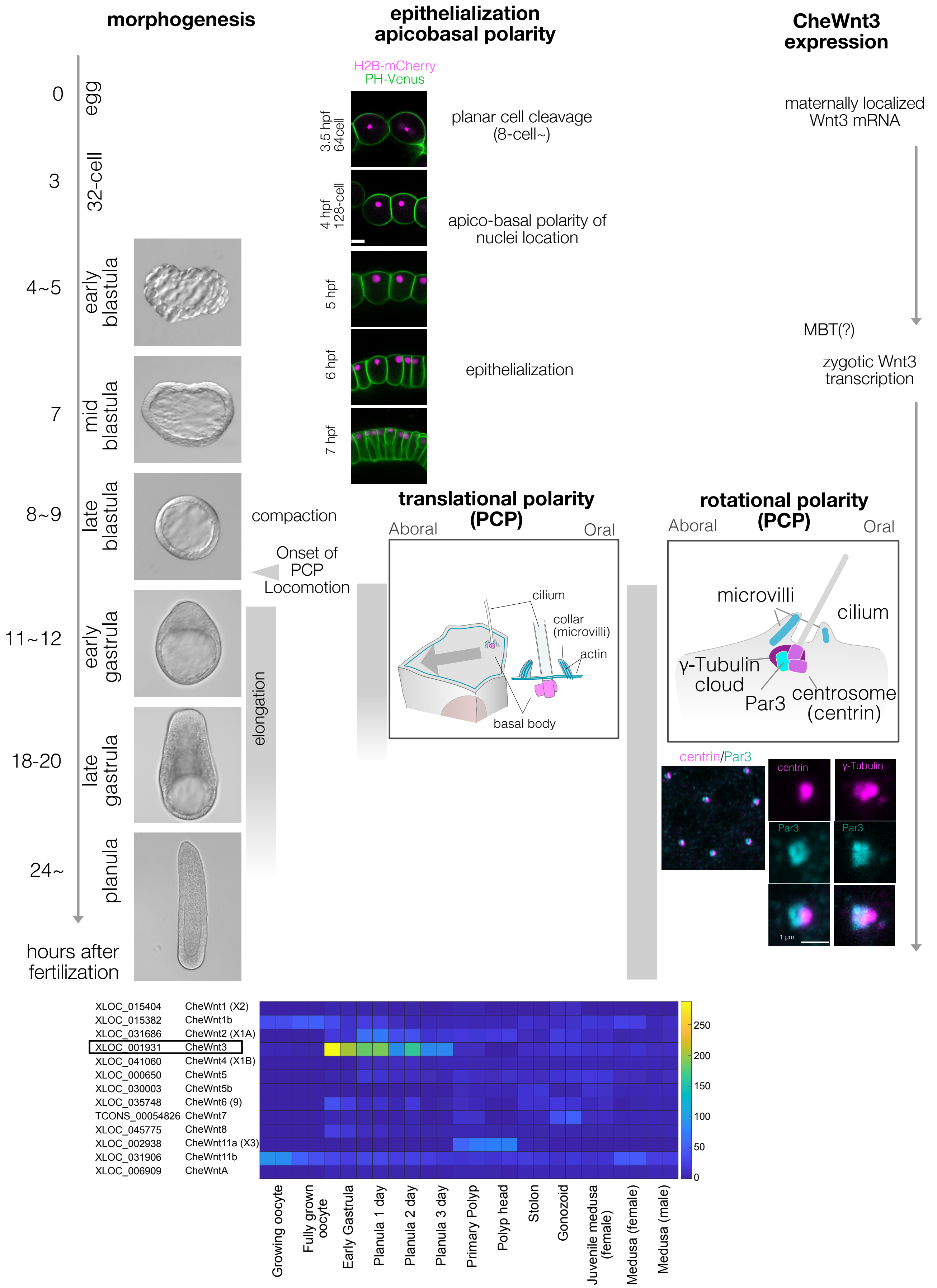
morphogenesis
epithelialization apicobasal polarity
CheWnt3 expression
0
egg
3
32-cell
4~5
early blastula
7
mid blastula
8~9
late blastula
11~12
early gastrula
18-20
late gastrula
24~
planula
hours after fertilization
compaction
Onset of PCP Locomotion
elongation
H2B-mCherry
PH-Venus
3.5 hpf 64cell
4 hpf 128-cell
5 hpf
6 hpf
7 hpf
planar cell cleavage (8-cell~)
apico-basal polarity of nuclei location
epithelialization
maternally localized Wnt3 mRNA
MBT(?)
zygotic Wnt3 transcription
translational polarity (PCP)
Aboral
Oral
cilium
collar (microvilli)
actin
basal body
rotational polarity (PCP)
Aboral
Oral
microvilli
cilium
γ-Tubulin cloud
Par3
centrosome (centrin)
centrin/Par3
centrin
γ-Tubulin
Par3
Par3
1 μm
XLOC_015404 CheWnt1 (X2)
XLOC_015382 CheWnt1b
XLOC_031686 CheWnt2 (X1A)
XLOC_001931 CheWnt3
XLOC_041060 CheWnt4 (X1B)
XLOC_000650 CheWnt5
XLOC_030003 CheWnt5b
XLOC_035748 CheWnt6 (9)
TCONS_00054826 CheWnt7
XLOC_045775 CheWnt8
XLOC_002938 CheWnt11a (X3)
XLOC_031906 CheWnt11b
XLOC_006909 CheWntA
250
200
150
100
50
0
Growing oocyte
Fully grown oocyte
Early Gastrula
Planula 1 day
Planula 2 day
Planula 3 day
Primary Polyp
Polyp head
Stolon
Gonozoid
Juvenile medusa (female)
Medusa (female)
Medusa (male)

**Figure S1 *Clytia* early embryogenesis stages and Wnt3 expression, related to Figure 1**

(A) Developmental stages of *Clytia hemisphaerica* early embryogenesis (18°C). The key events in morphogenesis, epithelialization and cell polarity are indicated. Morphologically, *Clytia* embryos form blastulae with monolayer blastomeres and blastocoel inside it. Clear blastocoel is developed from 16- or 32-cell stages (3~3.5 hpf). The morphological apico-basal polarity becomes prominent starting from the 128-cell stage, initially as apical localisation of the nuclei (4 hpf). The blastomeres epithelialize around the mid-blastula stage (7 hpf). The embryos become smaller (i.e. compaction) at the late blastula stage (8~9 hpf) when the PCP also start to be coordinated, and the embryo starts to swim with cilia beating. The PCP is almost fully coordinated by the early gastrula stage (11~12 hpf) when the OA-axis is morphologically prominent with the pointed-end shape at the oral end. The gastrulation occurs at the oral end by ingression of individual cells. The endodermal cells migrate from oral to aboral in the blastocoel. Body elongation along the OA-axis mainly occurs concomitantly with gastrulation and endoderm migration. Both the PCP and endoderm formation are required for the axial elongation. The PCP is initially prominent as the eccentric location of the ciliary basal body at the oral side of each cell (translational PCP) in the early gastrula stage. Cilium is surrounded by actin-rich microvilli collars. Starting from the late gastrula stage (18~20 hpf), the basal body structure and surrounding collar also become polarised along the OA-axis (rotational polarity); the F-actin positive collar is consistently reinforced at the aboral side, most likely mechanically supporting the directional power and recovery strokes along the OA axis. It is also the most reliable PCP marker in later stages (Momose 2012). The immunoreactivity of the Par3 (anti mouse Par3 antibody) was also detected at the aboral side of the ciliary body, suggesting a link between apico-basal cell polarity and PCP.

(B) Expression profile of *Clytia* Wnt genes in different developmental stages from existing RNAseq data (http://marimba.obs-vlfr.fr). A number of transcripts detected per million reads (TPM). The latest Wnt nomenclature based on Condamine et al. (DOI: 10.1016/j. ydbio.2019.09.001) and gene ID (XLOC) or transcript ID (TCONS) are indicated along with the previous names (Momose 2008) in parentheses.

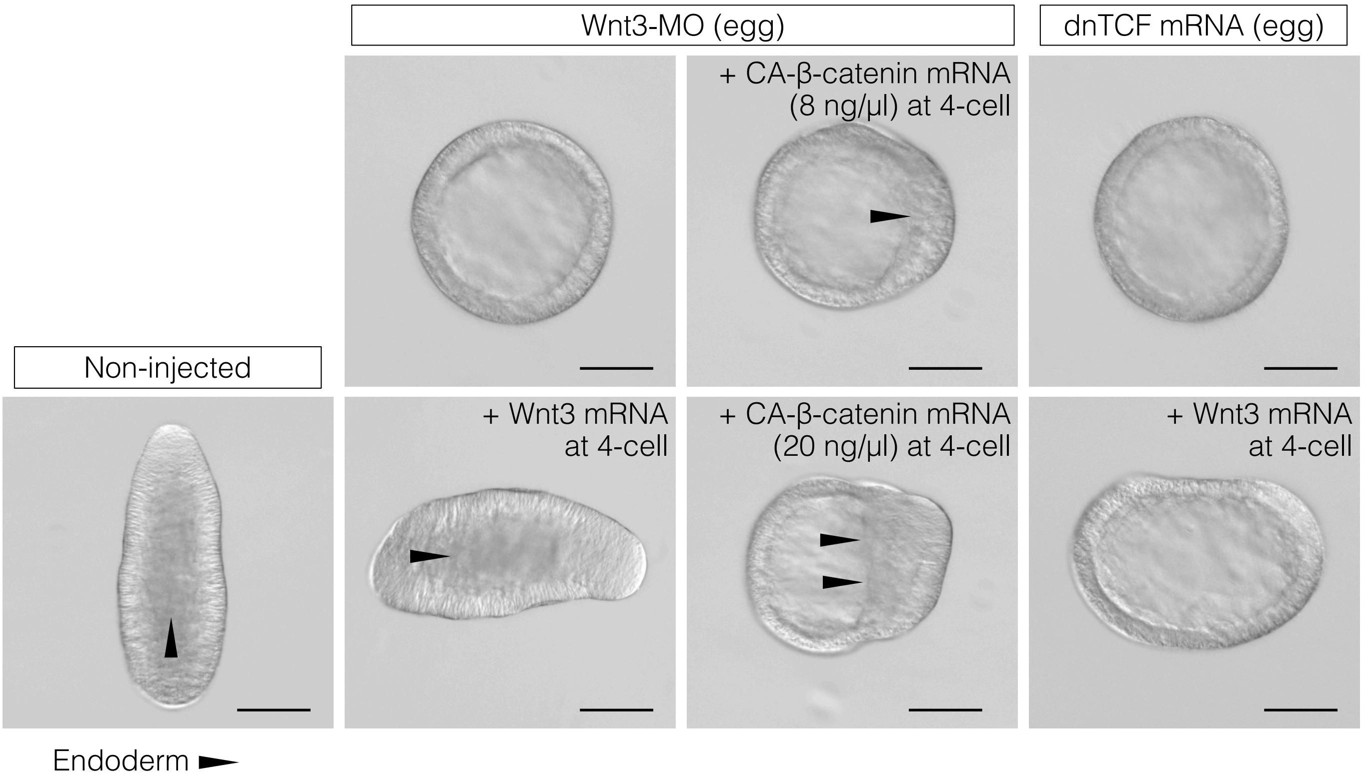


**Figure S2 CheWnt3 is necessary and sufficient to orient the morphological oral-aboral body axis, while the Wnt/β-catenin pathway acting under CheWnt3 is necessary but not sufficient. Related to Figure 3.** DIC observation of Non-injected (control) and Wnt3-MO or dominant negative form of CheTCF (dnTCF) mRNA was injected into the egg with an additional injection of CheWnt3 mRNA or constitutively active form of *Clytia* β-catenin (CA-β-catenin). The images are representative results used for the measurement in Fig. 3D. Bar= 100 µm. Triangles indicate the frontline of endoderm migration from the oral (Wnt/-β-catenin-activated side) toward the aboral.

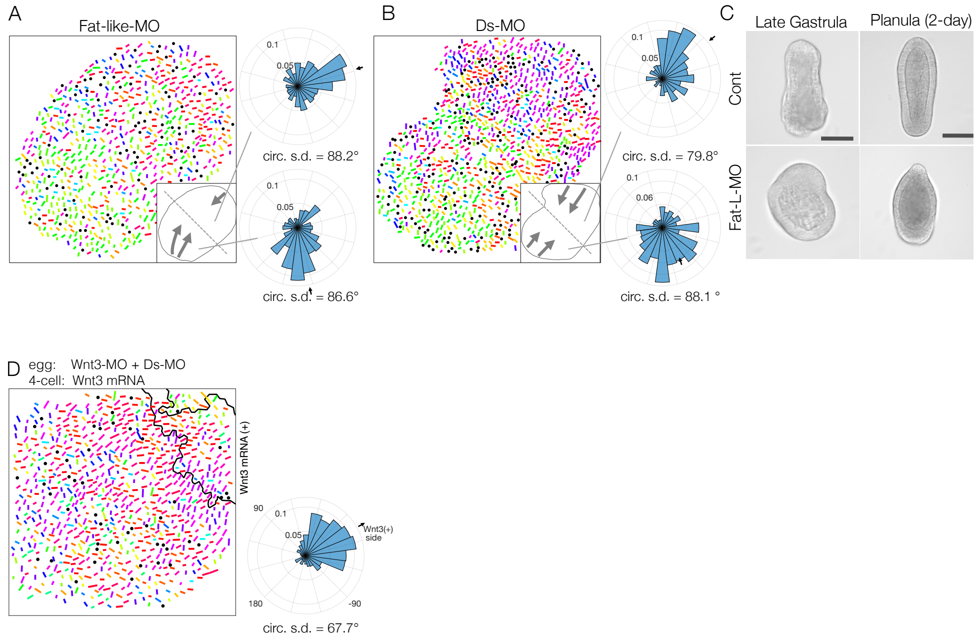


**Figure S3. Fat-like and Dachsous play roles in long-range PCP coordination for axial patterning in *Clytia* embryos. Related to Figure 4.**

(A,B) Long-range PCP coordination in Fat-like-MO injected early gastrula embryo (CheFat1, A) and Ds-MO injected embryo (CheDs, B). In both cases, while PCP is locally oriented to a specific direction, it is globally uncoordinated within an embryo. Two radar charts represent PCP orientation in halves of the single embryo. (C) Morphology of Fat-like-MO injected embryos at the late gastrula and 2-day planula stages. Axial elongation was severely impaired in Fat-like-MO injected embryos. Gastrulation from the oral end took place. The OA-axis polarity is present. (D) Wnt3-rescue by local mRNA injection can rescue the PCP defect by Ds-MO injection. CheWnt3 and Fat/Ds sysem may play partially overlapping roles for globally orienting PCP along the OA-xis xis morphologically remained distinctive, unlike Wnt3-MO or Stbm/Fmi-MO injected embryos. (D)

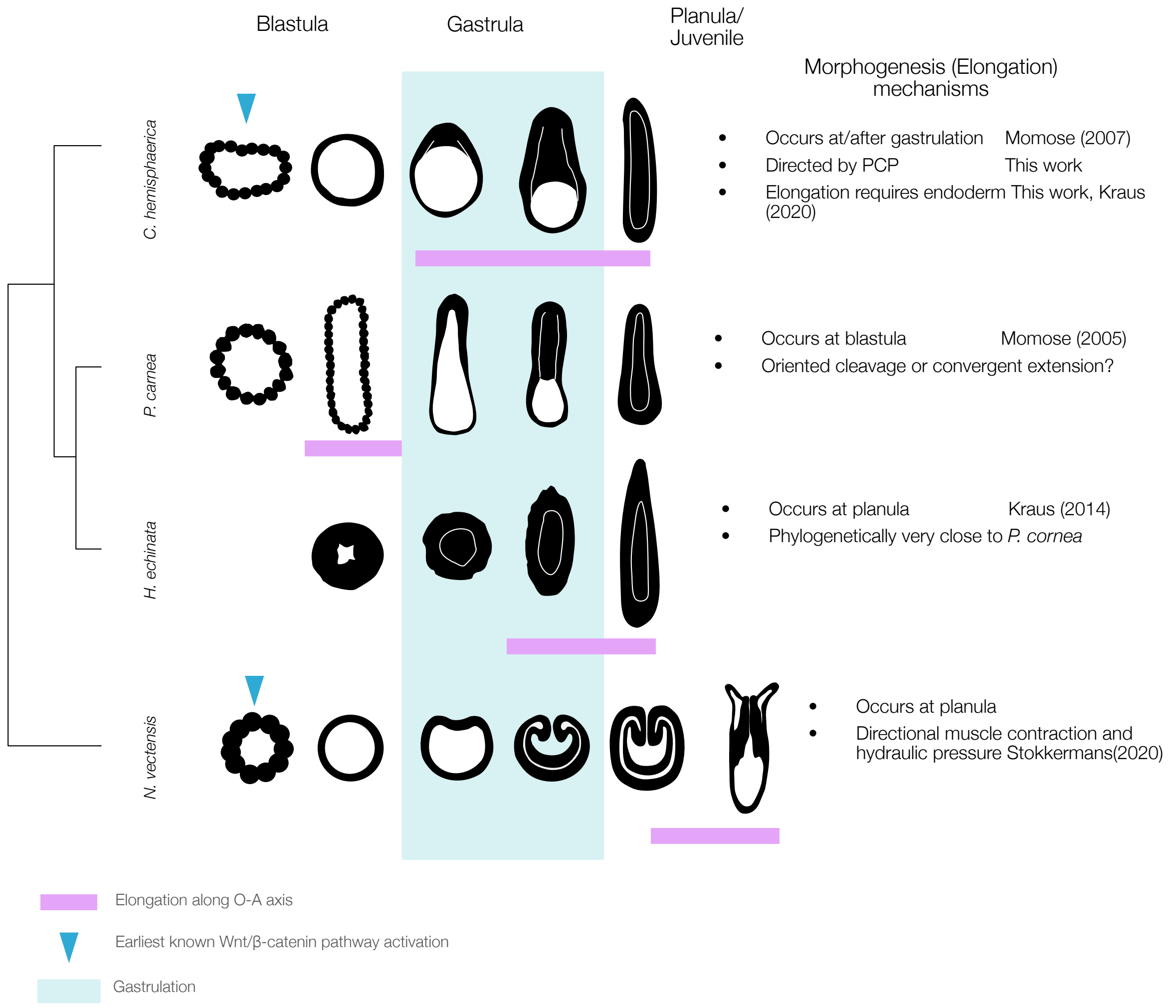


**Figure S4. A wide heterochronic variation of axial elongation in phylum Cnidarian.** The axial elongation process occurs at diffrernt timing (indicated by magenta box), along the key developmental stages in four cnidarian model species, *Clytia hemisphaerica*, *Podocoryne carnea*, *Hydractinia echinata* and *Nematostella vectensis*. The early gastrula stage is defined by the onset of the germ layer segregation and is typically the first stage, exhibiting a morphological oral-aboral axis. Gastrulation pattern is also highly variable; invagination, unipolar ingresssion, delamination from stereoblastula. The elongation occurs concomitantly to the gastrulation in *C. hemisphaerica*, while it occurs before the gastrulation in *P. carnea.* In contrast, in *N. vectensis*, elongation is mechanically driven by hydraulics caused by directional muscle contraction coordinated along the OA axis (Stokkermans et al., 2022). The elongation is thus driven by the existing tissue polarity. The role of the core PCP proteins in muscle PCP orientation remains to be studied in *N. vectensis*.

#### Figure S5

| | | | β-catenin | Axin | GSK3-β | TCF | Wnt | Fzd/Frizzled | Dsh/Dishevelled | Flamingo/Celsr | Strabismus/Vangl | Inversin | Prickle | Fat/Fat-like | Dachsous/Ds | PCP effectors |
|---|---|---|---|---|---|---|---|---|---|---|---|---|---|---|---|---|
| | | | Wnt/β-catenin | | | | | Core PCP | | | | | | Fat/Ds/Fj | | |
| Amoeba | | *Dictyostelium* | ±[15a] | | +[16] | | | | | | | | | | | |
| Fungi | | *Allomyces macrogynus* | | | | | | | | | | | | | | + |
| Filasterea | | *Capsaspora owczarzaki* | | | | | | | | | | | | | | + |
| Choanoflagellate | | *Salpingoeca rosetta* | ±[d] | ± | ++ | ± [b] | | | ± | -[1] | -[1] | +[1, 10] | +[1,10e] | ±[17f] | | + |
| | | *Monosiga brevicollis* | ±[d] | ± | ++ | ± [c] | -[9] | -[9] | ± | -[17] | -[17] | +[1, 10] | +[1,10e] | ±[17f] | | + |
| Ctenophora | | *Mnemiopsis leidyi* | +[8] | -[8] | +[8] | +[8] | +[8] | +[8] | +[8] | -[1] | -[1] | -[1] | +[1e] | | | + |
| Porifera | Homoscleromorpha | *Oscarella carmela* | +* | +[6] | +* | +* | +[6] | +[6] | +[6] | +[1] | +[1] | +[1] | +[1e] | ±[17f] | | + |
| | | *Oscarella lablaris* | +* | ? | +[5] | ? | +[5] | +* | +* | +[1] | +[1] | +[1] | +[1e] | | | + |
| | Calcerea | *Sycon ciliatum* | +[7] | -* | +[7] | +[7] | +[7] | +[7] | +[7] | +[1] | +[1] | +[1] | +[1e] | | | + |
| | Demospongiae | *Amphimedon queenslandica* | +[3,4] | +[3] | +[3,4] | +[3,4] | +[3,4] | +[3,4] | +[3,4] | +[1] | -[1] | +[1] | +[1e] | +[14,17] | | + |
| | Hexactinellida | *Aphrocallistes vastus* | +[2] | +[2] | +[2] | +[2] | -[2] | +[2] | -[2] | -[1] | -[1] | +[1] | +[1e] | | | + |
| Cnidaria | Anthozoa | *Nematostella* | + | + | + | + | + | + | + | +[1] | +[1] | +[1] | +[1e] | +[12] | +[12] | + |
| | Hydrozoa | *Clytia hemisphaerica* | + | + | + | + | + | + | + | +[1] | +[1] | +[1] | +[1] | +[12 g] | +[12] | + |
| | | *Hydra* | + | + | + | + | + | + | + | +[1] | +[1] | +[1] | +[1] | +[12g] | +[12] | + |
| Placozoa | | *Tricoplax adherans* | +[11] | +[11] | +[11] | +[11] | +[11] | +[11] | +[11] | +[1] | +[1] | -[1] | +[1] | +[13] | +[13] | + |
| Bilaterians | | | + | + | + | + | + | + | + | + | + | + | + | + | + | + |

**Figure S5. Both Wnt//β-catenin and core PCP genes are older than the sponge-cnidarian-bilaterian last common ancestor. Related to Figure 4.**

(A) Meta-analysis of the presence and absence of Wnt/β-catenin and core PCP protein genes in basal metazoan and non-metazoan species from existing works. Homoscleromorpha sponges *Oscarella* have a complete set of core PCP proteins. PCP controlled by core PCP proteins may be innovated in a sponge-cnidarian-bilaterian common ancestor, equivalent to the metazoan common ancestor in the sponge-ancestor scenario. The lack of PCP effectors in Fungi and *Capsaspora* and Inversin and Prickle orthologues in Choanoflagellate suggest primitive PCP regulatory mechanisms predate the metazoan ancestor.+: orthologue identified with fully conserved domains -: orthologue absent ±: a gene with homology sequence exists but lacks complete structure (domains). * identified in this work.

a: β-catenin in *Dictyostelium* is β-catenin-like aardvark, which is a component of the junctional complex and involved in cell signalling (not mediated by GSK3).
b: TCF-like in choanoflagellate has HMG domain but not TCF orthologue.
c: Dsh-like gene in choanoflagellida is a PDZ domain-containing protein lacking DEP/DIX domains.
d: Choanoflagellate β -catenin: Armagiro repeat-containing proteins.
e: Prickle in Ctenophoores//Spongs are Prickle-Testin common ancestor. Choanoflagelleate has Testin orthologue but not
f: Monosiga and Amphimedon lefftyrin cadherins share EGF and lamG domains with FAT cadherins, but their location (N-terminal) is different (adjacent to the transmembrane domain)
g: *Clytia* and *Hydra* have Fat1 (Fat-like) but not Fat (Fat4)


1 Schenkelaars et al., (2016) https://doi.org/10.1186/s12862-016-0641-0
2 Schenkelaars et al., (2017) https://doi.org/10.1038/s41598-017-15557-5
3 Adamska et al., (2010) https://doi.org/10.1111/j.1525-142X.2010.00435.x
4 Windsor Reid et al., (2018) https://doi.org/10.1186/s12862-018-1118-0
5 Lapébie et al., (2009) https://doi.org/10.1371/journal.pone.0005823
6 Nichols et al., (2006) https://doi.org/10.1073/pnas.0604065103
7 Leininger et al., (2014) https://doi.org/10.1038/ncomms4905
8 Pang et al., (2010) https://doi.org/10.1186/2041-9139-1-10
9 King et al., (2008) https://doi.org/10.1038/nature06617
10 Lapébie et al., (2011) https://doi.org/10.1002/bies.201100023
11 Srivastava et al., (2008) https://doi.org/10.1038/nature07191
12 Brooun et al., (2020) https://doi.org/10.1073/pnas.1917570117
13 Hulpiau and van Roy (2011) https://doi.org/10.1093/molbev/msq233
14 Abedin and King (2008) https://doi.org/10.1126/science.1151084
15 Grimson et al., (2000) https://doi.org/10.1038/35047099

### Table S1 Database accession numbers for Wnt/β-catenin and PCP genes. Related to Figure 4 and Figure S4

| | | | | Wnt/β-catenin | | | | | PCP (core PCP, Ft/Ds) | | | | | | | | |
|---|---|---|---|---|---|---|---|---|---|---|---|---|---|---|---|---|---|
| | | | | β-catenin | Axin | GSK3 | TCF | Wnt | Frizzled/Fzd | Dshevelled /Dvl | Flamingo /Celsr | Strabismus Van Gogh | Inversin /Diego | Prickle (Testin) | Fat/Fat-like | Dachsous | Source |
| Amoeba | | *Dictyostelium* | Yes | ±a | Absent | L34674 | Absent | Absent | Absent | Absent | Absent | Absent | Absent | Absent | Absent | Absent | Ensemble Protists |
| Fungi | | *Allomyces macrogynus* | Yes | ±d KNE54313 | Absent | KNE66930 | Absent | Absent | Absent | Absent | Absent | Absent | Absent | Absent | Absent | Absent | Ensemble Fungi |
| Filasterea | | *Capsaspora owczarzaki* | Yes | Absent | Absent | KJE89342 | Absent | Absent | Absent | Absent | Absent | Absent | Absent | Absent | Absent | Absent | Ensemble Protists |
| Choanoflagellate | | *Salpingoeca rosetta* | Yes | ±d XM_004991040 | Absent | XM_ 004990934 | ± b XM_ 004997546 | Absent | Absent | ±c | Absent (ref 1) | Absent (ref 1) | XM_004991658 | XM_ 004988894 | Absent (protocadherin lacking LamG) | ? | Genbank |
| | | *Monosiga brevicollis* | Yes | Absent | Absent | XM_ 001745535 | ± b XM_ 001749981 | Absent (ref 9) | Absent (ref 9) | HM448818.1 | Absent (ref 17) | Absent (ref 17) | XM_001746371 | XM_ 001742763 | ± f XM_ 001750799 | ? | Genbank |
| Ctenophora | | *Mnemiopsis leidyi* | Yes | ML073715a(HM448817) | Absent (ref 8) | ML20837a | ML11223a(HM448822) | ML01051a (HM448816), ML101013a (HM448815), ML06021a (HM448814), ML07528a (HM448813) | ML003224a (HM448819), ML034649b (HM448820) | ML005358a(HM448818) | Absent (ref 1) | Absent (ref 1) | Absent (ref 1) | ML34571a(ML2189) | ? | ? | Ensemble Metazoa |
| Porifera | Homoscleromorpha | *Oscarella carmela* | Yes | EB741523 | comp23540_c0_seq2 | comp23762_c0_seq1, comp23762_c0_seq3 | comp28653_c0_seq3 | comp8600_c0_seq1, comp49400_c0_seq1, comp36292_c0_seq12, comp30539_c0_seq3, comp36292_c0_seq5, comp24555_c0_seq2, comp24555_c0_seq1, comp15540_c0_seq2, comp27463_c0_seq2, comp28793_c0_seq3, comp27694_c0_seq2, comp35766_c0_seq3, comp15540_c0_seq1, comp27694_c0_seq5 | AJE25507-AJE25509 | comp44599_c0_seq1 | comp27089_c0_seq1 | comp31109_c0_seq5 | comp36587_c0_seq1, comp33916_c0_seq2 | comp26691_c0_seq1 | ? | ? | Compagen Japan, Genbank |
| | | *Oscarella lobularis* | No | GQ144649 | ? | GQ144648 | ? | GQ144646, GQ144647 | KM365029, KM365030, KM365031 | KT898949 | KT898950 | KT898956 | KT898953-KT898954 | KT898955 | ? | ? | Genbank |
| | Demospongiae | *Amphimedon queenslandica* | Yes | NM_001279305.1 | XM_ 019993695.1 | NM_ 001279283 | HM561507.1 | GQ144651 ABX90060 GQ144650 | NP_001266206, NP_001266202 | NP_ 001266242 | Aqu1.224307 | Absent (ref1) | Aqu1.221107, Aqu1.227557 | Aqu1.217773 | XM_ 011408514 | ? | Genbank |
| | Hexactinellida | *Aphrocallistes vastus* | Yes | comp17679_c0_seq1 | comp13763_c0_seq1 | comp17686_c0_seq3 | comp24352_c0_seq1 | Absent (ref 2) | comp96115_c0_seq1 | Absent (ref 2) | Absent (ref 1) | Absent (ref 1) | comp15586_c0_seq1, comp34481_c0_seq1 | comp27734_c0_seq1 | ? | ? | Compagen |
| Cnidaria | Anthozoa | *Nematostella vectensis* | Yes | XM_001647467 | XM_ 048720266 | XM_ 001628384 | DQ497247 | XP_001641494, XP_ 001633773, XP_001635899, XP_001623100, XP_ 001630693, XP_001629451, XP_001630717, XP_ 001630717, XP_001630032, XP_001637576, XP_ 001629450, XP_001629964, XP_001632885, XP_ 001637670 | XM_001647490, XM_032379260, XM_048734489 | XM_ 032380552 | XM_ 032365472 | XM_001630370 | XM_032362457 | XM_ 032367105 | XM_ 032383036 (Fat1), XM_ 048724363.1 (Fat4) | XM_ 048733391 | Genbank |
| | Hydrozoa | *Clytia hemisphaerica* | Yes | XLOC_003560 (JQ438997) | XLOC_ 011804 | XLOC_016339 (EU374723) | XLOC_ 007658 (JQ806376) | XLOC_15404, XLOC_15382, XLOC_31686, XLOC_01931, XLOC_41060, XLOC_00650, XLOC_30003, XLOC_35748, XLOC_34538, XLOC_45775, XLOC_02938, XLOC_06909 | XLOC_044260 (DQ869571) XLOC_039886 (JQ439004) XLOC_010791 (DQ869572) XLOC_002646 (JQ439005) | XLOC_ 015396 (JQ439000) | XLOC_ 001639 (JQ439002) | XLOC_013328 (JQ439008) | XLOC_037832 (JQ438998) | XLOC_010268 (JQ439007) | XLOC_ 033731 (partial JQ439001) | XLOC_ 037142 (JQ438999) | Marimba (Genbank) |
| | | *Hydra vlugaris* | Yes | U38624 | XM_ 002155907 | NM_ 001309721 | NM_ 001309733 | AB426122, AB426123, MH940211, AB425832, AB426121, AB426120, AB425835, AB425836, AB426119, AB425834, AB425833, | NM_001309725, NM_001280959, XM_002163078 | NM_ 001287794 | XM_ 047290573 | XM_047273357 | XM_012701755 | XM_ 047270537 | XM_ 012702814 | XM_ 047270896 | Genbank |
| Placozoa | | *Tricoplax adherans* | Yes | XM_002110267 | XM_ 002109742 | XM_ 002111683 | XM_ 002116135 (partial) | XM_002116071, XM_ 002108187 | AJE25511, AJE25512 | No accession | XM_ 002118173 | XM_002107607 | Absent (ref 1) | XM_ 002112366 | XM_ 002116901 | XM_ 002112915 | Genbank |
| Bilaterians | | | | + | + | + | + | + | + | + | + | + | + | + | + | + | |

#### Figure S6

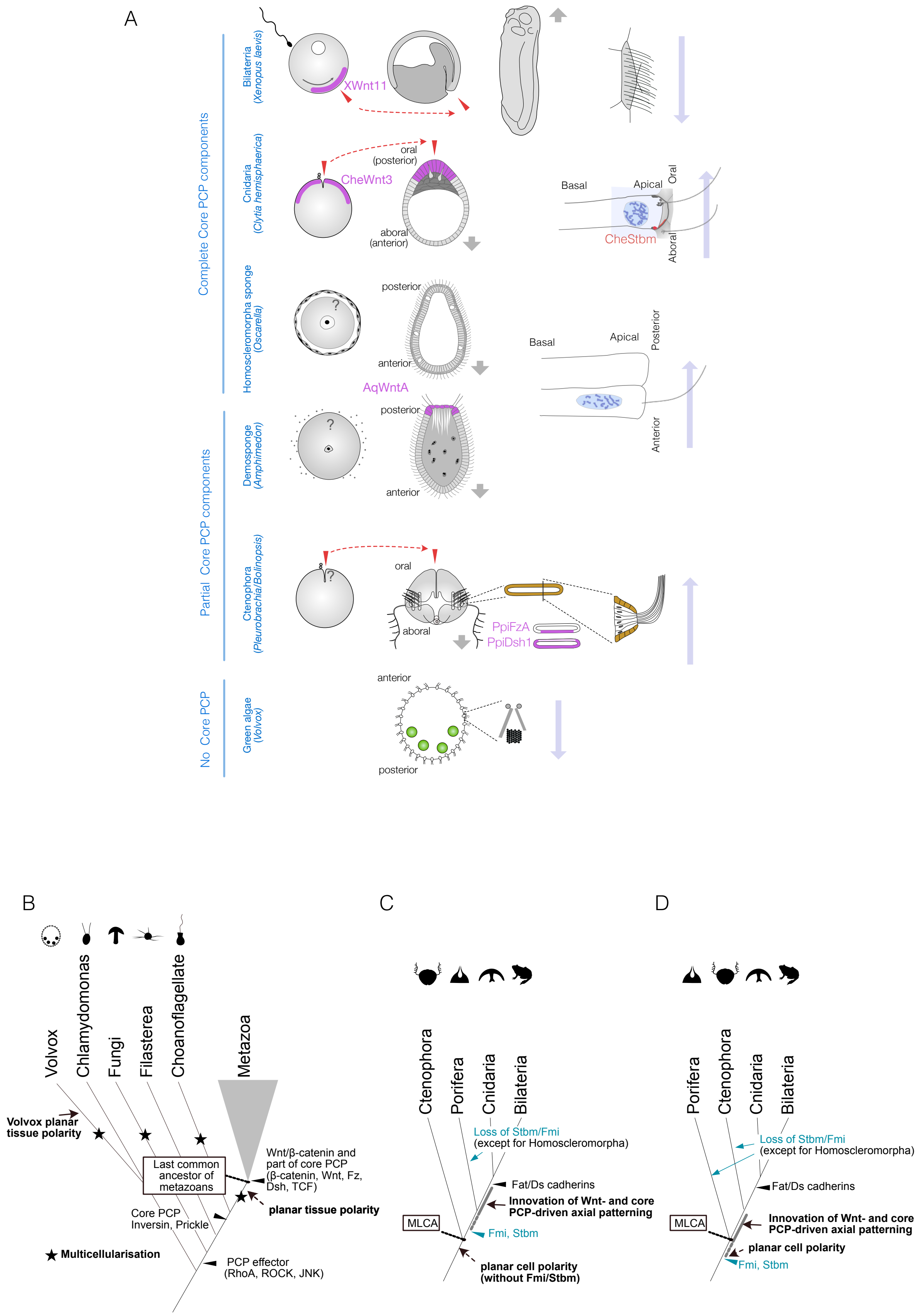
A
Complete Core PCP components
Partial Core PCP components
No Core PCP
Bilateria (Xenopus laevis)
XWnt11
Cnidaria (Clytia hemisphaerica)
oral (posterior)
CheWnt3
aboral (anterior)
Basal
Apical
Oral
Aboral
CheStbm
Homoscleromorpha sponge (Oscarella)
posterior
anterior
AqWntA
Basal
Apical
Posterior
Anterior
Demosponge (Amphimedon)
posterior
anterior
Ctenophora (Pleurobrachia/Bolinopsis)
oral
aboral
PpiFzA
PpiDsh1
Green algae (Volvox)
anterior
posterior
B
Volvox
Chlamydomonas
Fungi
Filasterea
Choanoflagellate
Metazoa
Volvox planar tissue polarity
Last common ancestor of metazoans
Wnt/β-catenin and part of core PCP (β-catenin, Wnt, Fz, Dsh, TCF)
planar tissue polarity
Core PCP Inversin, Prickle
Multicellularisation
PCP effector (RhoA, ROCK, JNK)
C
Ctenophora
Porifera
Cnidaria
Bilateria
Loss of Stbm/Fmi (except for Homoscleromorpha)
Fat/Ds cadherins
Innovation of Wnt- and core PCP-driven axial patterning
MLCA
Fmi, Stbm
planar cell polarity (without Fmi/Stbm)
D
Porifera
Ctenophora
Cnidaria
Bilateria
Loss of Stbm/Fmi (except for Homoscleromorpha)
Fat/Ds cadherins
MLCA
Innovation of Wnt- and core PCP-driven axial patterning
planar cell polarity
Fmi, Stbm

**Figure S6. PCP, or coordinated polarity of motile cilia, is tightly coupled to axial patterning across metazoans and is independently evolved in non-metazoan species. Related to Figure.4**

(A) Planar cell/tissue polarity associated with the body axis in metazoan and multicellular green algae *Volvox*. Egg polarity and embryonic axial anatomy are indicated on the left. Cell polarity associated with the body axis is indicated on the right. The localisation of Wnt ligand or other localised PCP factors associated with egg polarity or body axis is indicated with a purple shade. In *Clytia*, *Xenopus* and ctenophores, egg polarity determines the embryonic body axis, and these coordinations are shown in pairs of red arrows. The locomotion and fluid flow created by motile cilia are indicated with grey and long blue arrows, respectively. *Volvox*, a multicellular green algae which independently acquired multicellularity and has no Wnt or core PCP proteins, has clearly coordinated planar tissue polarity. The PCP thus can occur without Wnt and core PCP proteins. (B-D) Evolutionary scenarios of the innovation of the body axis and PCP. (B) The events in the common ancestor of the metazoan and its outgroups. Multicellrisation events that occurred multiple times are indicated with an asterisk (★). Key components of Wnt/β-catenin and a part of core PCP pathways (β-catenin, Wnt ligand, Frizzled, Disheveled, TCF) predates the last common ancestor of metazoans (MLCA). A part of the core PCP proteins, Inversin(Diego)/Testin and Prickle are innovated in metazoan and choanoflagellate common ancestor and all PCP effectors (Rho GTPase; RhoA, cdc42, Rock etc) regulating cytoskeletal arrangements are older or as old as the last common ancestor of fungi and metazoans (opisthokonts), from Figure S5. (C) Metazoan evolution with ctenophore-ancestor scenario and (D) sponge-ancestor scenario. In either case, the full set of the core PCP protein has evolved in the sponge-cnidaria-bilaterian common ancestor. Stbm/Fmi was lost multiple times in sponges, which should have occurred in the ctenophore in the sponge-ancestor scenario. Wnt ligand and Dishvelled are also lost in Hexactinellida *Aphrocallistes vastus* and *Oopsacas minuta* (Francis et al. https://doi.org/10.1098/rsos.230423 and Schenkelaars et al. https://doi.org/10.1038/s41598-017-15557-5)
